# Development of a large-scale computer-controlled ozone inhalation exposure system for rodents

**DOI:** 10.1101/489518

**Authors:** Gregory J. Smith, Leon Walsh, Mark Higuchi, Samir N. P. Kelada

**Affiliations:** Department of Genetics, Curriculum in Toxicology & Environmental Medicine, University of North Carolina at Chapel Hill, Chapel Hill, NC, USA; United States Environmental Protection Agency, Research Triangle Park, NC, USA

**Keywords:** ozone, air pollution, inhalation, exposure, computer-controlled, exposure system

## Abstract

Complete systems for laboratory-based inhalation toxicology studies are typically not commercially available; therefore, inhalation toxicologists utilize custom-made exposure systems. Here we report on the design, construction, testing, operation, and maintenance of a newly developed *in vivo* rodent ozone inhalation exposure system. Key design requirements for the system included large-capacity exposure chambers to facilitate studies with large sample sizes, automatic and precise control of chamber ozone concentrations, as well as automated data collection on airflow and environmental conditions. The exposure system contains two Hazelton H-1000 stainless steel and glass exposure chambers, each providing capacity for up to 180 mice or 96 rats. We developed an empirically tuned proportional-integral-derivative (PID) control loop that provides stable ozone concentrations throughout the exposure period (typically 3h), after a short ramp time (∼8 minutes), and across a tested concentration range of 0.2 to 2 ppm. Specific details on the combination analog and digital input/output system for environmental data acquisition, control, and safety systems are provided, and we outline the steps involved in maintenance and calibration of the system. We show that the exposure system produces consistent ozone exposures both within and across experiments, as evidenced by low coefficients of variation in chamber ozone concentration and consistent biological responses (airway inflammation) in mice, respectively. Thus, we have created a large and robust ozone exposure system, facilitating future studies on the health effects of ozone in rodents.

## Introduction

Ozone (O_3_) exposure is associated with significant short and long-term adverse health effects (Levy et al. 2001; Ito et al. 2005; Levy et al. 2005; Jerrett et al. 2009). As a powerful oxidant and respiratory irritant, O_3_ reacts with airway lining fluid constituents to generate biologically active compounds that cause immediate lung function decrements and acute toxicity (Pryor et al. 1995; Nielsen et al. 1999; Mudway and Kelly 2000). Within hours of inhalation exposure to O_3_, there is an inflammatory response in the respiratory tract characterized by cytokine release and an influx of macrophages and neutrophils in the lung (Aris et al. 1993; Devlin et al. 1996; Mudway and Kelly 2000). Downstream consequences of O_3_ exposure include increased susceptibility to respiratory infections and exacerbations of existing airway diseases such as asthma and chronic obstructive airway disease (Ko et al. 2007; Ji et al. 2011; Kim et al. 2011). Long-term exposure to O_3_ has also been linked to increased incidence of diabetes, cardiovascular disease, neurological disease, and mortality (Turner et al. 2016; Day et al. 2017; Jerrett et al. 2017; Cleary et al. 2018).

Although its general biological effects are well studied, the toxicological mechanisms underlying these adverse health effects of O_3_ remain the subject of ongoing research. Particularly active areas of research include, but are not limited to, identification of specific biologically active products of airway surface liquid ozonation (Speen et al. 2016), the mechanisms of systemic cardiovascular and neurological effects (Paffett et al. 2015; Miller, Snow, et al. 2016; Miller, Ghio, et al. 2016; Tyler et al. 2018), and the influence of O_3_ on the pathogenesis (vs. exacerbation) of respiratory diseases (Herring et al. 2015; Michaudel et al. 2018; Zu et al. 2018). Additionally, the mechanisms underlying inter-individual differences in responses to ozone due to sex (Cabello et al. 2015; Cho et al. 2018 Sep 21) and genetic variation (Bauer and Kleeberger 2010) are also important data gaps that need to be addressed. A combination of epidemiological and experimental approaches (controlled *in vivo* human and rodent studies, and *in vitro* systems) will be needed to answer all of these questions. We have focused our work on the use of rodent models to study ozone toxicity, including inter-individual differences in response. In particular, we aim to identify genetic predictors of ozone response using a population of genetically diverse mice, which requires quite large sample sizes (e.g., more than 300 mice). To facilitate such large-scale inhalation studies, we designed and constructed a new large-scale, computer-controlled O_3_ exposure system for rodents at the University of North Carolina.

Inhalation exposure systems typically require a great deal of custom engineering. The design of exposure systems require the expertise of inhalation toxicologists and engineers, and individual components often need to be acquired from several different commercial sources or fabricated inhouse. Because we found these aspects to be true in developing our exposure system, we have written this manuscript to serve as a helpful reference for others considering developing a similar exposure system. The general design as well as many of the components used for the construction of our chambers will likely work well for systems of different sizes implemented by other investigators.

There are several reviews describing guidelines on the best practices for inhalation toxicology studies (Phalen 1976; Dorato 1990; Pauluhn 2003; Wong 2007; Chen and Lippmann 2015). Detailed manuscripts on the design of inhalation exposure systems for gases such as O_3_ and various other classes of airborne toxicants such as particulate matter, aerosols, and vapors have also been published (O’Shaughnessy et al. 2003; Wong 2007; McKinney and Frazer 2008; Goldsmith et al. 2011). Our goal was to design and implement an ozone exposure system that complies with published guidelines (Phalen 1976; Chen and Lippmann 2015; OECD 2018), while at the same time meeting our specific experimental requirements. General design considerations for whole-body inhalation toxicology studies include a delivery system for clean air, compatible/inert construction materials, control and characterization of the exposure atmosphere, and maintenance of suitable environmental conditions without the buildup of waste gases (Phalen 1976). In addition to these general requirements, we specifically needed the ability to expose a large number of mice to minimize the number of batches (which could confound analyses), an automated control system to maintain precise and reproducible exposures, and automated monitoring and recording of data on several environmental parameters. Here we report on the design, construction and performance of a large-capacity exposure system for rodents that features automatic and precise control of chamber ozone concentrations, as well as automated data collection on airflow and environmental conditions.

## Materials and Methods

### Exposure System Design and Operation

Block diagrams of the complete system are shown in Figures 1A and B and screen captures from the computer control system are shown in Figures 2A and B. For reference, major individual components used to construct the system are provided in Table 1 and detailed wiring diagrams are provided as Supplemental Data.

**Figure 1.**
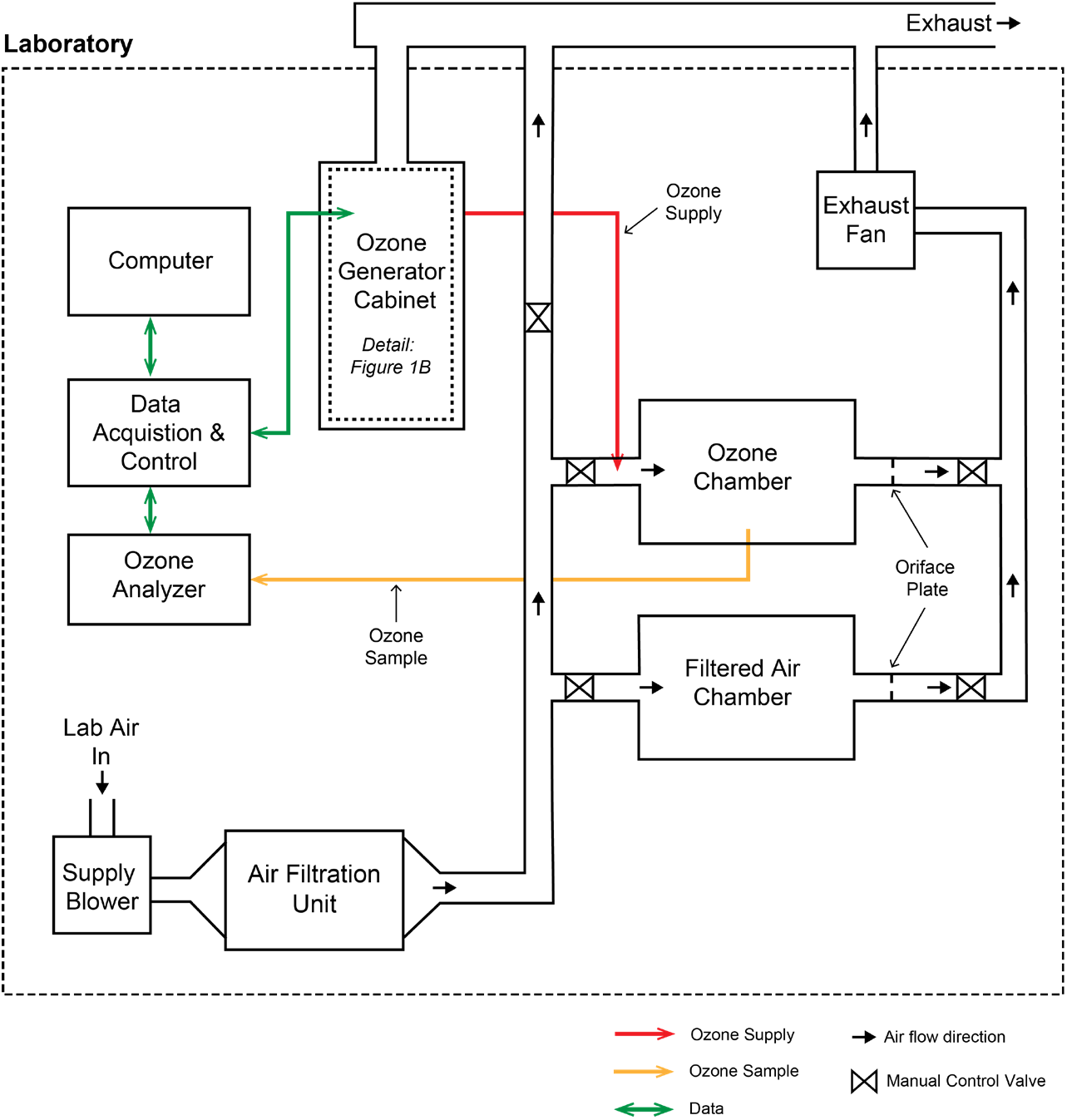
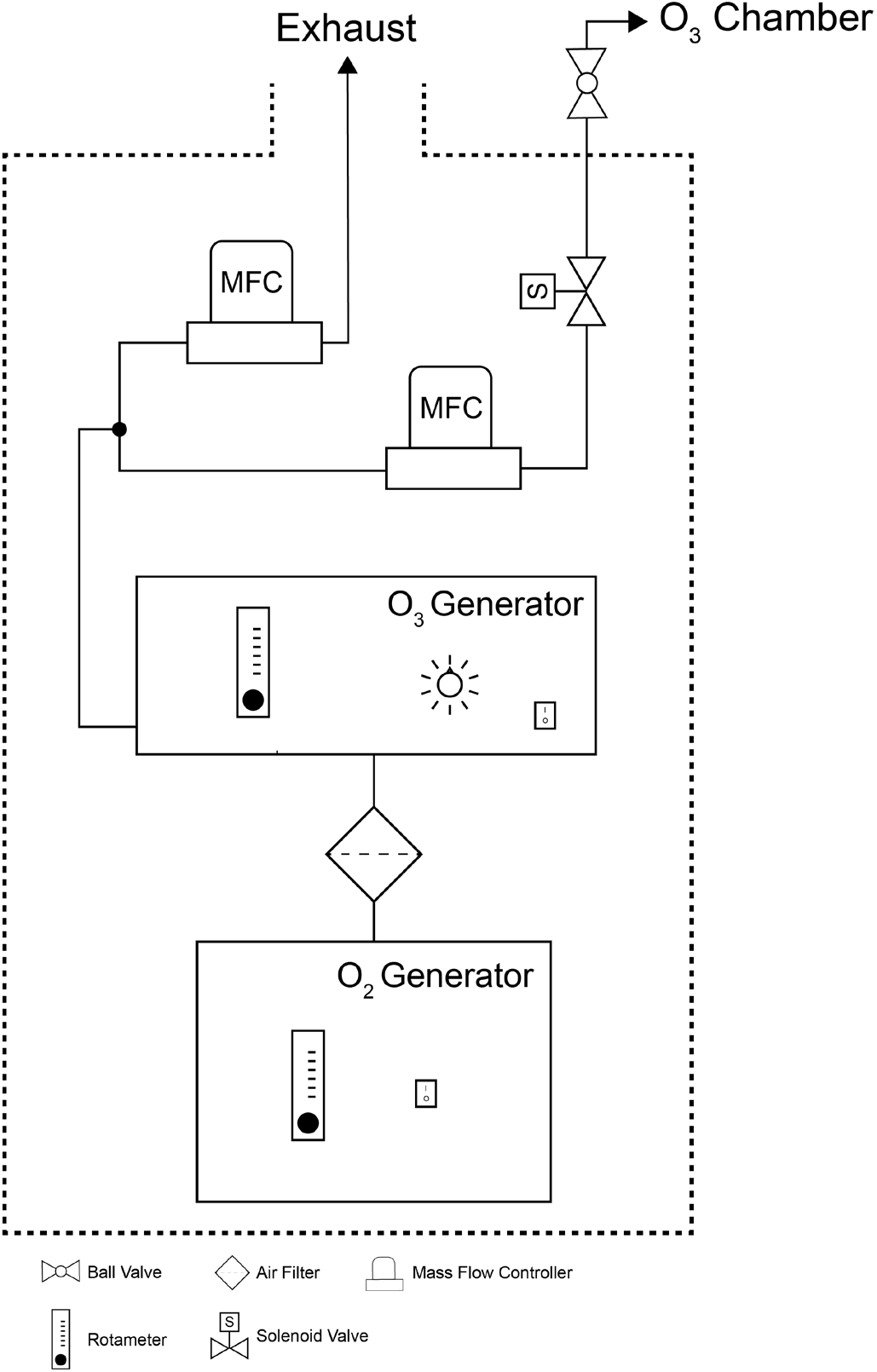
Block diagrams of the exposure system. (A) Overview of major components including the clean air supply, chambers, ozone control system, chamber and cabinet exhaust. (B) Detail of O_2_ and O_3_ generator cabinet. Compressed, concentrated oxygen flows from the oxygen generator to the ozone generator then enters a stainless steel manifold attached to the supply and waste mass flow controllers. The waste flow vents into the cabinet exhaust. Downstream of the supply mass flow controller within the gas safety cabinet is a relay actuated solenoid valve. Outside the cabinet is a manual control valve before the O_3_ supply tubing enters the chamber supply pipe.

**Figure 2.**
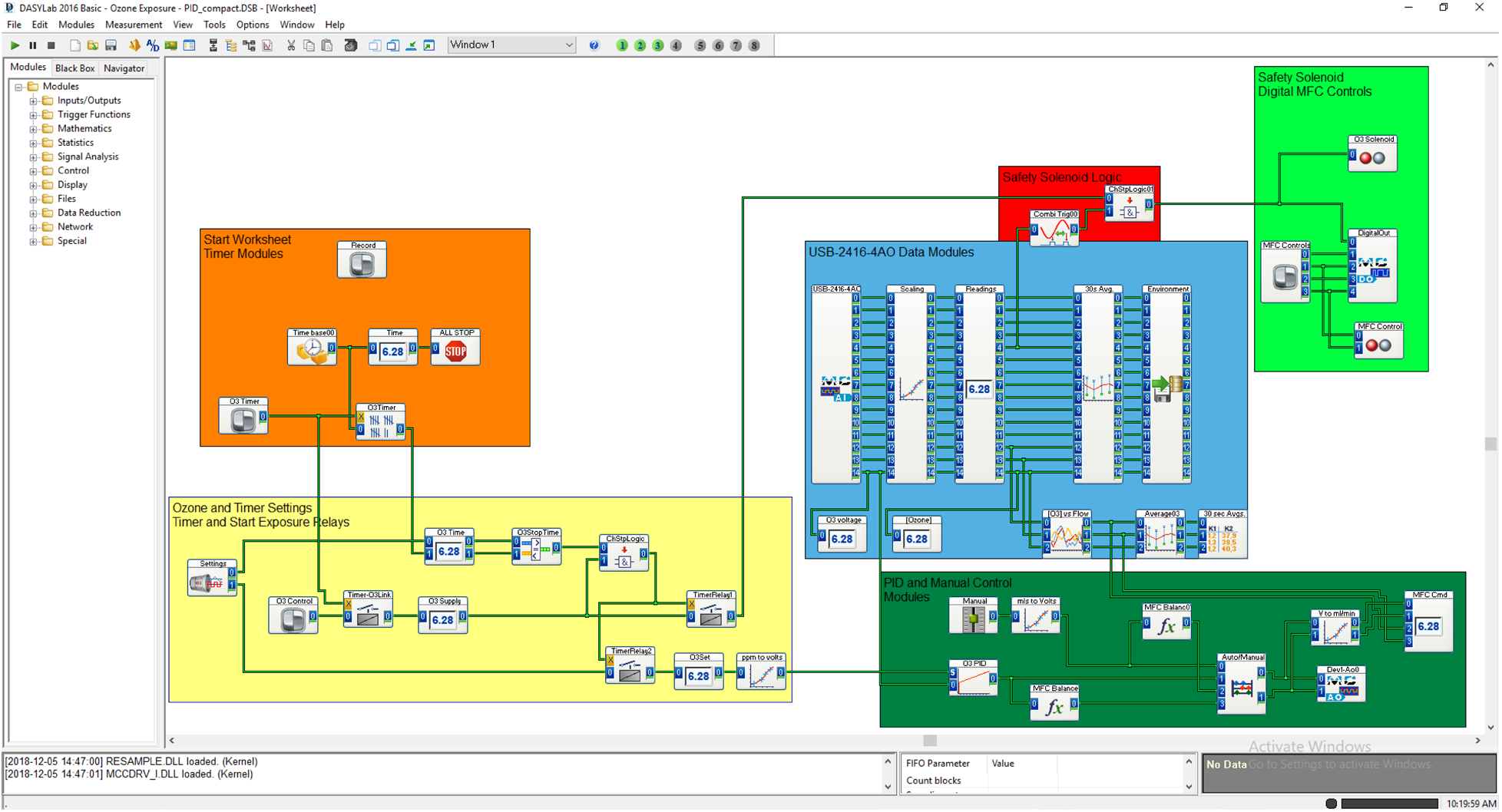
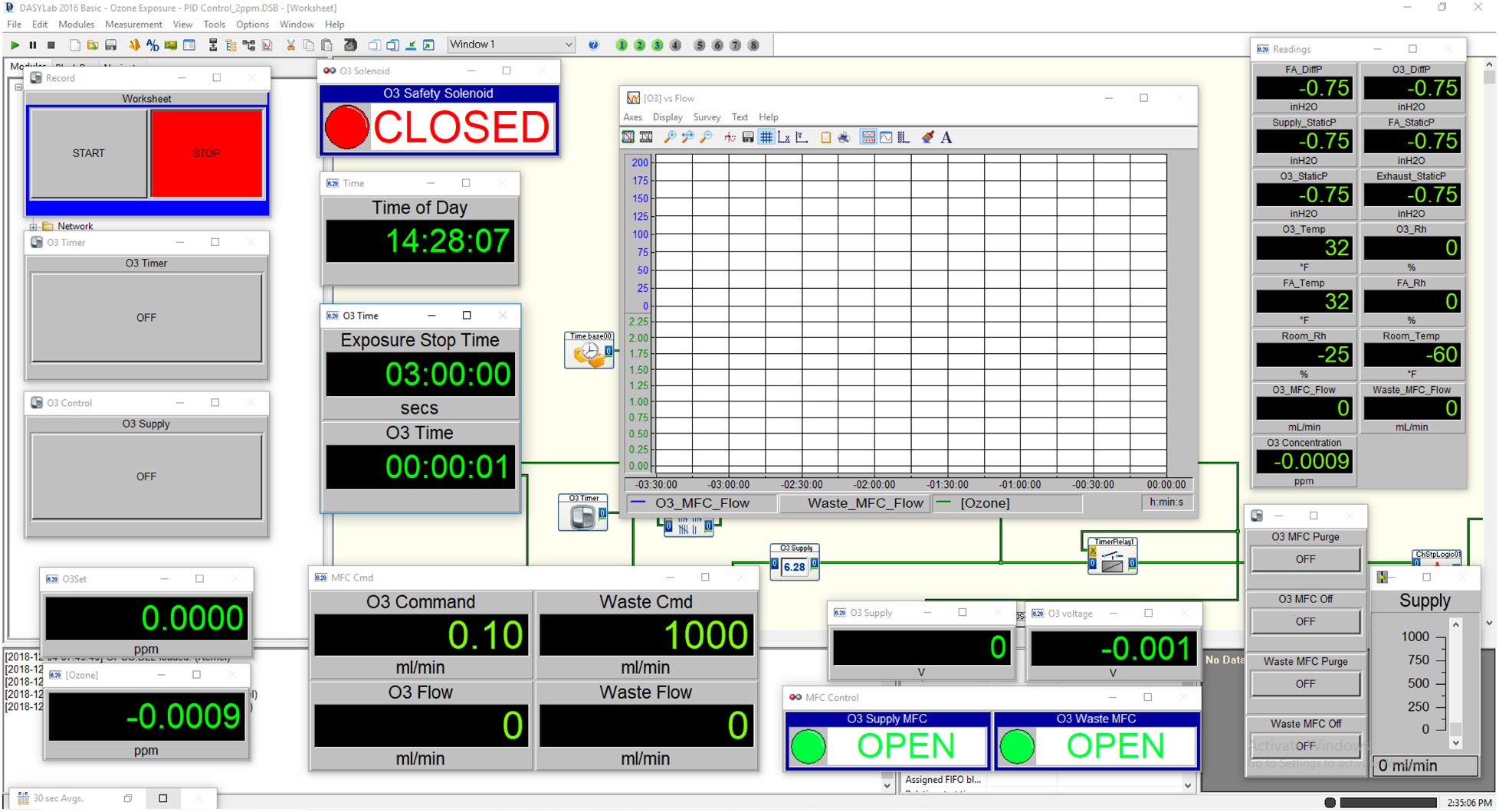
Screen captures of the DASYLab^®^ computer-control system. (A) Layout of worksheet modules used to acquire and record environmental data, control the MFCs (manual and automatic), operate the safety shutoff program, and control the duration and concentration of exposures. (B) View of worksheet showing switches, digital displays, chart recorder, manual control slider, and valve status indicators.

**Table 1.** List of major exposure system components.

| **System Component** | **Product Description** | **Manufacturer** | **Location** |
| --- | --- | --- | --- |
| **Pressure Blower** | Model FPB-70F-M5 | Greenheck | Schofield, WI |
| **Chamber Ozone Analyzer** | Model 49i Ozone Analyzer | Thermo Scientific | Waltham, MA |
| **Chamber T/Rh Transmitters** | HX94BV1W | Omega | Norwalk, CT |
| **Computer** | Optiplex 3040 | Dell | Round Rock, TX |
| **Control Software** | DASYLab® 2016 Basic V. 14.0 | Measurement Computing | Norton, MA |
| **Custom Air Filtration Unit Housing** | 4-Stage Side Access | AAF Flanders | Louisville, KY |
| **Exhaust Fan** | Model FR110 | Fantech | Lenexa, KS |
| **Exposure Chamber** | Hazelton H-1000 | Lab Products | Seaford, DE |
| **Gas Safety Cabinet** | 7200 2-Cylinder | Safety Equipment Corporation | Belmont, CA |
| **Laboratory Ozone Detector** | C-30ZX | EcoSensors | Newark, CA |
| **Mass Flow Controllers** | Aalborg GFC-17 | Aalborg | Orangeburg, NY |
| **Oxygen Generator** | Onyx A-016 | A2Z Ozone Inc. | Louisville, KY |
| **Ozone Generator** | 4G Lab Benchtop | A2Z Ozone Inc. | Louisville, KY |
| **Pressure Transmitters** | DM-2006-LCD | Dwyer | Michigan City, IN |
| **Room T/Rh Transmitter** | RHP-2W11-LCD | Dwyer | Michigan City, IN |
| **USB Data Acquisition and Control** | USB-2416-4AO | Measurement Computing | Norton, MA |

*T= Temperature*
*Rh= Relative humidity*

### Exposure chambers

We designed the system to provide controlled, filtered air and air plus ozone exposure atmospheres using two commercially available Hazelton H-1000 stainless steel and glass whole-body inhalation exposure chambers (Lab Products)(Brown and Moss 1981). Two doors at the front and rear of the chambers provide access to the animal cages and visual monitoring of animals in the chamber. Within the chambers, three levels of wire mesh cage racks are mounted on sliding tracks with pans to collect excreta beneath each cage rack. Cage racks accommodate up to 60 mice each (larger cages are available for rats and other laboratory animals). The cage racks feature removable food trays and an automatic watering system to facilitate animal housing and/or multiple days of exposure. The H-1000 chambers have several ports in each glass door and in the top of the steel chamber body. We used three ports on each chamber to monitor: 1) static air pressure relative to the laboratory, 2) ozone concentration, and 3) temperature and relative humidity. The top and bottom of the chambers have 3-inch diameter clamp-style sanitary fittings that we connected to the stainless steel air supply ducts and CPVC exhaust piping, respectively.

### Air supply and exhaust system

At the top of the chambers, we connected a filtered air supply system. Filtered, temperature, and humidity conditioned laboratory air, supplied by a pressure blower, is passed through a custom air filtration unit made by AAF Flanders (AAF, Smithfield, NC). The filtration unit, consisting of a side access filter housing and 4 stages of filters (Stage 1: 2” Minimum Efficiency Reporting Value (MERV) 8 prefilter, Stage 2: 12”-SAAF^TM^ gas filtration cassette, Stage 3: MERV 8 prepost filter, Stage 4: MERV 14 Final filter) removes ambient particulate matter, ozone, and other gases from the air stream. Following the filtration unit, air enters a 3-inch diameter stainless steel ducted supply manifold. The supply manifold divides the filtered air into an excess supply vent (to exhaust) and two chamber drops. We installed manual control valves at the excess supply vent and chamber drops to adjust supply manifold pressure and chamber inlet airflow, respectively. Immediately downstream of each of the chambers are 3-inch CPVC exhaust lines with orifice plates and individual flow control valves (chamber flow and static pressure balance) that combine at an exhaust manifold and then pass through an inline duct fan to enter the laboratory’s exhaust system.

### Chamber balancing/airflow calibration

We designed the system to have a nominal airflow rate of 250 L/min (15-air changes/hour/chamber) and a slight negative chamber pressure relative to the room to minimize the possibility of ozone entering the laboratory. We achieved these parameters by sequentially adjusting the supply and exhaust fan speeds and the system of control valves. While adjusting the system of fans and valves, we measured the static and differential pressures at specific locations of the air supply, chambers, and exhaust. First, the operating pressure ranges of the chamber supply and exhaust manifolds were determined by running the supply blower and exhaust fans while the valves to the chambers were closed. After we achieved a sufficient difference between the supply and exhaust manifold static pressures (relative to the laboratory), we opened the chamber valves and adjusted them until the measurement of an orifice flow meter installed in the chamber exhaust corresponded to 250 L/min. The orifice flow meter we constructed consists of a flow restrictive orifice plate and a pressure transmitter connected to taps in the exhaust pipe to measure the differential pressure between up and downstream of the orifice. To generate a calibration curve for the orifice flow meter, we made airflow measurements with a pitot tube placed several feet downstream of the orifice and plotted these measurements against differential pressure measurements at a series of exhaust and supply valve settings. The calibration curve enables the maintenance of airflow at 250 L/min based on the corresponding orifice flow meter reading. To enable the monitoring of pressure readings directly (without the computer control software open), we mounted all of the static and differential pressure transmitters (6 units) in the door of a wall cabinet where they are visible. We also installed electrical conduit with wires to carry 4-20mA current signals from the pressure transmitters to the opposite side of the laboratory. We terminated the wires at a USB microcontroller, which converts the 4-20mA analog signal to a digital signal enabling visualization and recording on a computer.

### Ozone Generation and Analysis

Ozone generation equipment is housed in a compressed gas safety cabinet (Figure 1B). The cabinet has an air vent on the front door and an exhaust on the top of the cabinet, which is connected to the laboratory’s exhaust ventilation and maintains a negative pressure inside the cabinet to prevent leaks. A medical oxygen concentrator was chosen for the system due to its safety, and lower cost over time compared to compressed oxygen. The oxygen generator (A2Z Ozone, Louisville, KY) was placed in the lower section of the cabinet, and connected with plastic tubing through an in-line particle filter to the ozone generator. We maintained the flow of oxygen to the ozone generator at the manufacturer’s recommended flow of 2 L/min. At this flow, the ozone generator is capable of generating up to 5 grams/hour of ozone via silent corona arc discharge. Downstream of the generator, we maintained a constant 1L/min flow of ozone by using the generator’s built-in rotameter. Ozonated air flows through fluorinated ethylene propylene (Teflon FEP) tubing into a 316 stainless steel manifold, which we connected to two Aalborg GFC17 mass flow controllers (MFC), designated supply and waste. The MFCs control the flow of ozone into the chamber (supply) and meter any excess ozone (waste) into the laboratory exhaust via the gas cylinder cabinet vent. The ozone supply MFC and the waste MFC are adjusted proportionally via an analog voltage signal (0-5 vDC) from a PC-based data-acquisition and control system. Downstream of the ozone supply MFC, we installed a relay-actuated solenoid safety valve and a manually operated cutoff valve. Following the control valves, we connected the ozone supply line to the stainless steel filtered air supply duct via a threaded compression fitting. This fitting was welded into the duct in between the control valve and the designated ozone chamber’s air inlet on the chamber drop duct. A stem extends into the center of duct that opens counter flow into the oncoming filtered air stream, which aids in mixing the ozone and filtered air streams as they enter the chamber. The H-1000 chambers have a circular plate centered below the chamber inlet that aids in uniformly distributing the incoming air or air-ozone mixture. We measure chamber ozone concentration with a Thermo 49i UV photometric ozone analyzer, which continuously draws chamber air at a flow rate of 3 liters/min through 1/4 inch FEP Teflon tubing inserted into the middle of the front door of the ozone chamber through a compression fitting. The sampling tube extends several inches into the chamber above the middle cage rack. The Thermo 49i analyzer then determines the chamber ozone concentration every 10 seconds. For personnel safety, we mounted an EcoSensors (Newark, CA) C30-ZX ozone monitor with an audible alarm and an LED light-bar graph display in the laboratory.

### Environmental Data Acquisition

Data from the ozone analyzer and environmental sensors (pressure, temperature, relative humidity, and chamber airflow) are continuously transmitted to a multifunction USB-based 24-bit data acquisition device (MCC-DAQ 2416-4AO) via a 4-20mA or 0 to 5vDC signal then displayed and recorded (30 second averages) by a computer. The MCC-DAQ 2416-4AO features 16 analog voltage input channels, to which the transmitter signals are passed. Additionally, the interface hardware included digital in/out (DIO) and 4 0-5 vDC analog out (AO) channels For transmitters that output an analog current of 4-20 mA, we converted the signal to a 0-5vDC voltage with a 250-ohm resistor placed in the circuit. The pressure transmitters have a measurement range of 0 to 3 inches of H_2_O, corresponding to a signal output of 4-20 mA. To allow for measurements of both positive and negative pressure, the transmitters have high and low-pressure connections. To measure positive pressure (e.g. supply manifold), we use the high-pressure connection and the low-pressure connection is open to the laboratory air. For low-pressure measurements (e.g. exhaust manifold or chamber static), the opposite is true. Differential pressure measurements for airflow required the use of both connections. We monitor the chamber temperature and relative humidity using combination temperature and relative humidity sensors inserted into ports on the rear door of each chamber. We also installed a temperature and relative humidity transmitter on a wall in the laboratory. The Thermo 49i analyzer has analog voltage outputs, which we used to transmit the ozone concentration signal to the USB microcontroller. Serial port or Ethernet based TCP/IP connections were also available for monitoring data from the Thermo 49i instrument.

### Computer-Control System

We developed a data acquisition and control system using a Windows-based PC that we can operate from within the exposure facility or remotely from our main lab via a Windows remote desktop connection. The system utilizes DASYLab^®^, an icon-based software that allows users to quickly develop customized data acquisition, analysis, and control applications without needing to write code. We employed a single worksheet in DASYLab^®^ containing a series of icons/modules to facilitate the acquisition and recording of environmental data, control of the MFCs (manual and automatic), operation of a safety shutoff program, and control the duration of exposures (Figure 2). We used DASYLab^®^ modules to scale voltage measurements from each analog input channel (e.g. 0 to 5 vDC = 0 to 5ppm ozone), display real-time data onscreen, calculate averages for data reduction, and record the data in a text file. We copied specific data streams and passed them to other modules to achieve four main objectives: 1) monitor and automatically control chamber ozone concentration at a desired set point during exposures, 2) control the duration of exposure, 3) provide a safety routine to automatically shut off the flow of ozone, and 4) monitor, display, and record all chamber environmental condition data. For initial chamber testing, we controlled the flow of ozone manually using slider modules connected to analog voltage outputs (MFC control signal). For automatic control during exposures, we used a proportional-integral-derivative (PID) control module in the DASYLab^®^ worksheet. We passed an analog voltage input from the ozone analyzer and a virtual-voltage from a signal generator module corresponding to the desired setpoint (e.g. 2 ppm setpoint = 2 vDC output) to the PID module. The PID module operates by continuously determining the deviation between the ozone analyzer voltage and the setpoint voltage to calculate an appropriate control signal output to the ozone supply MFC to minimize the deviation. We adjusted the gain settings (proportional, integral, derivative) and the maximum control output within the PID module to achieve an optimum exposure profile (Figure 3). We designed a series of modules that operate as a timer and a relay to control the duration of exposure and control the safety solenoid valve. Following the desired exposure time, the system turns off the ozone supply MFC output and closes the safety solenoid valve. We designed a safety routine using a trigger module to monitor the ozone chamber static pressure data stream and control a relay-operated, normally closed solenoid valve (safety solenoid valve). An increase in ozone chamber pressure indicating a leak or other problem with the air balance, will turn off the trigger module and thus the flow of ozone to the chambers. We set the trigger module so that it will only open the safety solenoid when both the static pressure is below a threshold setting (<0.10 inches H2O) and the timer-relay module is on.

**Figure 3.**
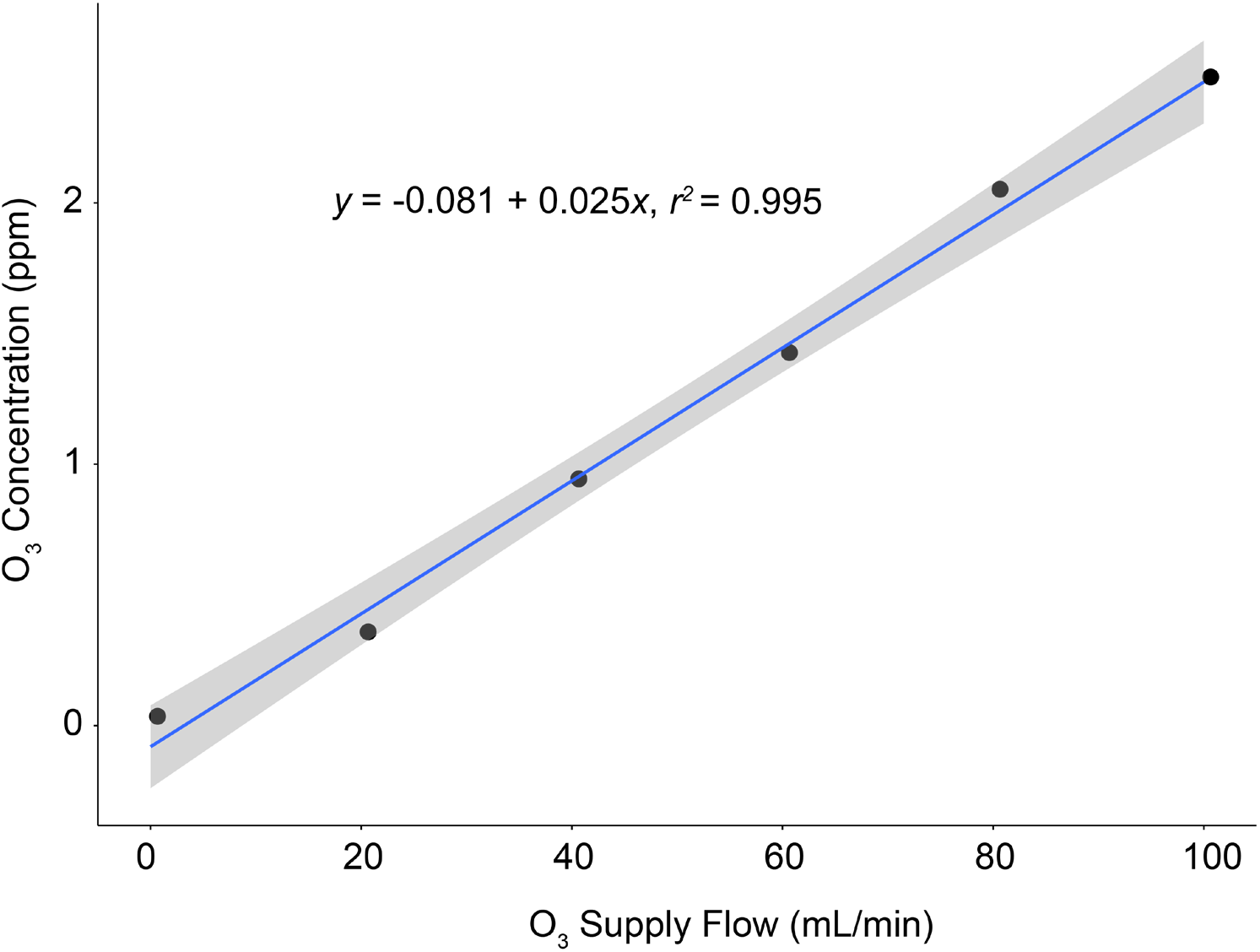
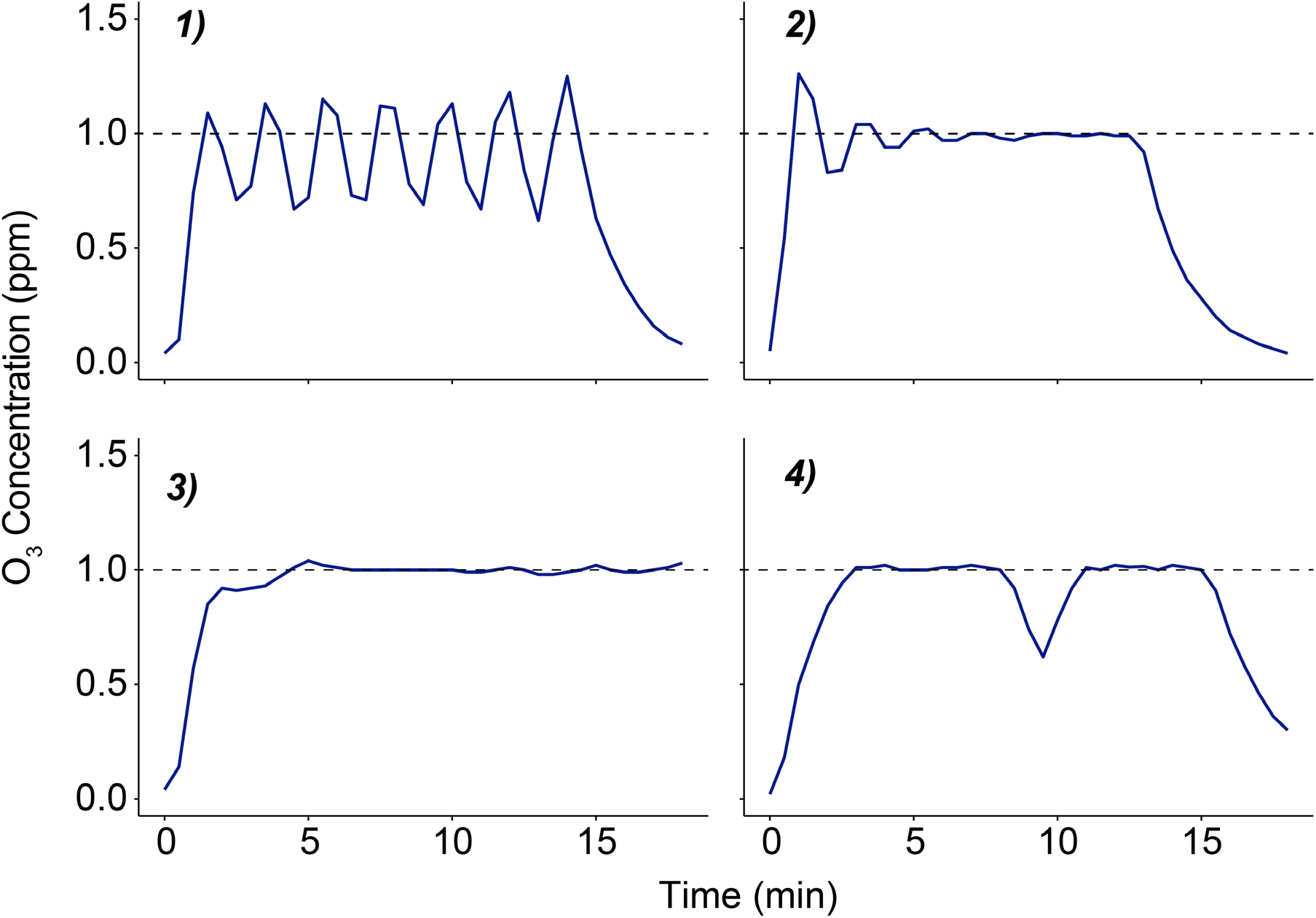
Initial manual testing and abridged PID control tuning. (A) Linear correlation between O_3_ supply MFC flow rate setting and resulting chamber O_3_ concentration. These data were used to determine the range and estimate the capacity of the generation system. The shaded area represents a 95% confidence interval for a linear regression model (see equation). (B) Panels illustrating the progression of the trial and error process used to determine optimal gain settings for the DASYLab^®^ PID control module. Panel 1: steady oscillations induced by increasing the proportional gain. Panel 2: improved settling time and offset resulting from increasing the integral and derivative gains. Panel 3: an appropriately damped system with optimal PID gain settings. Panel 4: system response to turning off the O_3_ generator for one minute. Dashed line indicates the desired concentration setpoint of 1 ppm during initial testing.

### Chamber Maintenance

While most of the components of the system are durable (e.g. chambers, valves, ducts, etc.) and require infrequent attention, three aspects of maintenance should be discussed, namely the air filtration unit, the ozone generation system, and ozone analyzer. The service life of the diluent air filtration stages will vary by frequency of use and ambient environmental conditions. For our system, we implemented general replacement schedules and guidelines for evaluating the function of the filtration unit. The MERV 8 filters are replaced every 30 exposure-hours or if noticeably soiled upon visual inspection (before every exposure). The MERV 14 filter has a longer service life and is replaced every 60 exposure-hours. We can estimate the function of the gas filtration cassette by observing the magnitude of the decrease in pre-exposure chamber ozone concentration from ambient levels. We plan to replace the cassette when it can no longer decrease the chamber ozone concentration to less than 20% of the ambient level prior to the start of an exposure. We monitor the function of the ozone generator and oxygen generator by the use of a handheld oxygen sensor, and observing the average level of flow required to maintain ozone concentrations at the set point. A significant departure from the oxygen generator’s typical output of 90% or in the average level of ozone supply flow compared to initial test exposures can indicate problems with the ozone generation system. The manufacturer’s manuals for both the ozone and oxygen generators contain troubleshooting guides in case there is a loss of function. We conduct zero and span checks prior to each exposure, using a zero air source and an ozonator within the instrument (the internal ozonator is an optional feature). Calibration of the ozone analyzer is conducted on a monthly basis with a Thermo 49i-PS transfer standard photometer. The transfer standard photometer is calibrated annually against the SRP-1 at the U.S. EPA Office of Research and Development in Research Triangle Park, NC.

### Concentration Uniformity Assay

To evaluate the concentration uniformity of ozone in the chamber, we utilized an assay developed by Flamm in which the oxidation of a 0.1M boric acid-buffered 1% potassium iodide indicator solution (BKI), measured by the absorbance of BKI at 352 nm in a UV-transparent 96 well plate, is used as a surrogate of ozone concentration (Flamm 1973). We distributed 35-mm polystyrene tissue culture dishes containing 3 milliliters of BKI throughout the chambers (five dishes in each of the three cage racks, one in each corner and one in the center). In the control chamber, we placed one dish in the middle of each rack. We conducted a 30-minute computer-controlled 2-ppm ozone exposure of the dishes. Finally, we took the standard deviation (SD) and coefficient of variability (CV) statistics of absorbance values across all of the dishes as an indicator of heterogeneity of the ozone concentration.

### Animal Chamber and Cage Sanitation

The H1000 chambers, trays, and cage racks were cleaned and sanitized prior to exposures. A solution of 70 percent ethanol was used to clean the inside surfaces of the H1000 animal chambers. Cage racks and trays were removed from the chambers and cleaned using a tunnel-style cage washer operated by the Department of Comparative Medicine at UNC.

### Animal Exposures and Phenotyping

We conducted animal exposure experiments using 8-10 week old female C57BL/6J mice obtained from The Jackson Laboratory. The mice were housed in an AAALAC approved facility and all procedures received approval by the Institutional Animal Care and Use Committee (IACUC) at the University of North Carolina Chapel Hill. Mice were housed over ALPHA-Dri bedding (Shepard) under standard 12h lighting conditions, with *ad libitum* food and water.

For exposures, we transferred the mice from their normal housing and placed them in individual stainless steel wire mesh cage racks within the H-1000 chambers. After the doors to the chambers were sealed, we started the automated ozone exposure worksheet and exposure timer module in DASYLab^®^. We exposed groups of mice to filtered air or 2-ppm ozone for 3 hours. After shut down, the chamber ozone concentration returned to ambient levels in approximately 10 minutes and we returned the mice to their normal housing. At 21 hours following exposure, we euthanized the mice by a lethal dose of urethane followed by exsanguination and bronchoalveolar lavage (BAL) fluid collection. We quantified percent neutrophils in the BAL fluid by standard differential white blood cell counting techniques.

## Results

### Chamber Testing and Operation

#### Manual Testing

Before performing tests with ozone present in the chambers, we evaluated the function of the safety shutoff routine. With the ozone generator unplugged and the rest of the system running, we turned the exhaust fan off or opened the chamber door. Both of these conditions caused the system to close the safety solenoid, confirming that the safety program would function if needed. To evaluate the performance of the exposure system and linearity of the relationship between ozone supply flow and chamber ozone concentration we conducted a manual step test (Figure 3A). With the PID module set point corresponding to 0 ppm ozone, we used a slider module to increase flow of ozone manually in steps of 20 ml/min. At each step, we allowed the chamber concentration to reach a plateau (at ∼8 minutes) then remain at steady state for at least 15 minutes before increasing the output for the next step. We concluded the test one step after reaching 2 ppm ozone, the maximum concentration planned for animal exposures in the lab. From the step test, we determined that we would need a flow rate of approximately 80 ml/min ozone to maintain a 2 ppm exposure.

#### Automated-Control Tuning and Exposure Profiles

Based on the results of the manual step test, we limited the maximum control output from the PID module to 0.5 vDC (100 ml/min) to prevent accidental overproduction of ozone during testing and provide some capacity (20 ml/min) to compensate for ozone loss due to uptake and adsorption by animals in the chamber. There are several published heuristic methods for determining, also known as “tuning”, the optimal gain settings of a PID controller (Astrom and Hägglund 1995; Arab and Mp 2012). While these references were helpful for understanding the basics of PID control theory, we ultimately determined the proportional, integral, and derivative gains for our PID control module through a trial and error process. An abridged visualization of the progress of tuning the PID control module is shown in Figure 3B. First, we adjusted the P, I, and D gain settings individually until we determined how each setting modulated the system’s response. Adjusting the P gain upward eventually produced a steady oscillation of the O_3_ concentration close to the desired set point (Figure 3B-1). Increasing the D gain caused the oscillations to taper off, and adding in an I-gain setting at this point eliminated a small offset causing the mean value to be below the set point (Figure 3B-2). We continued to increase the D-gain and decreased the P-gain slightly, which decreased the amount of settling time until the exposure system produced a fast-ramp and maintenance of chamber O_3_ at the setpoint with minimal overshoot (Figure 3B-3). These settings also produced a sufficient response to a brief loss of output from the O_3_ generator, returning the O_3_ concentration to the setpoint after we turned off the O_3_ generator for one minute (Figure 3B-4). The automated PID control loop was able to keep the ozone concentration to within 1% of the setpoint over several exposures, with and without mice in the chambers (Figure 3B-3, and Figure 4).

**Figure 4.**
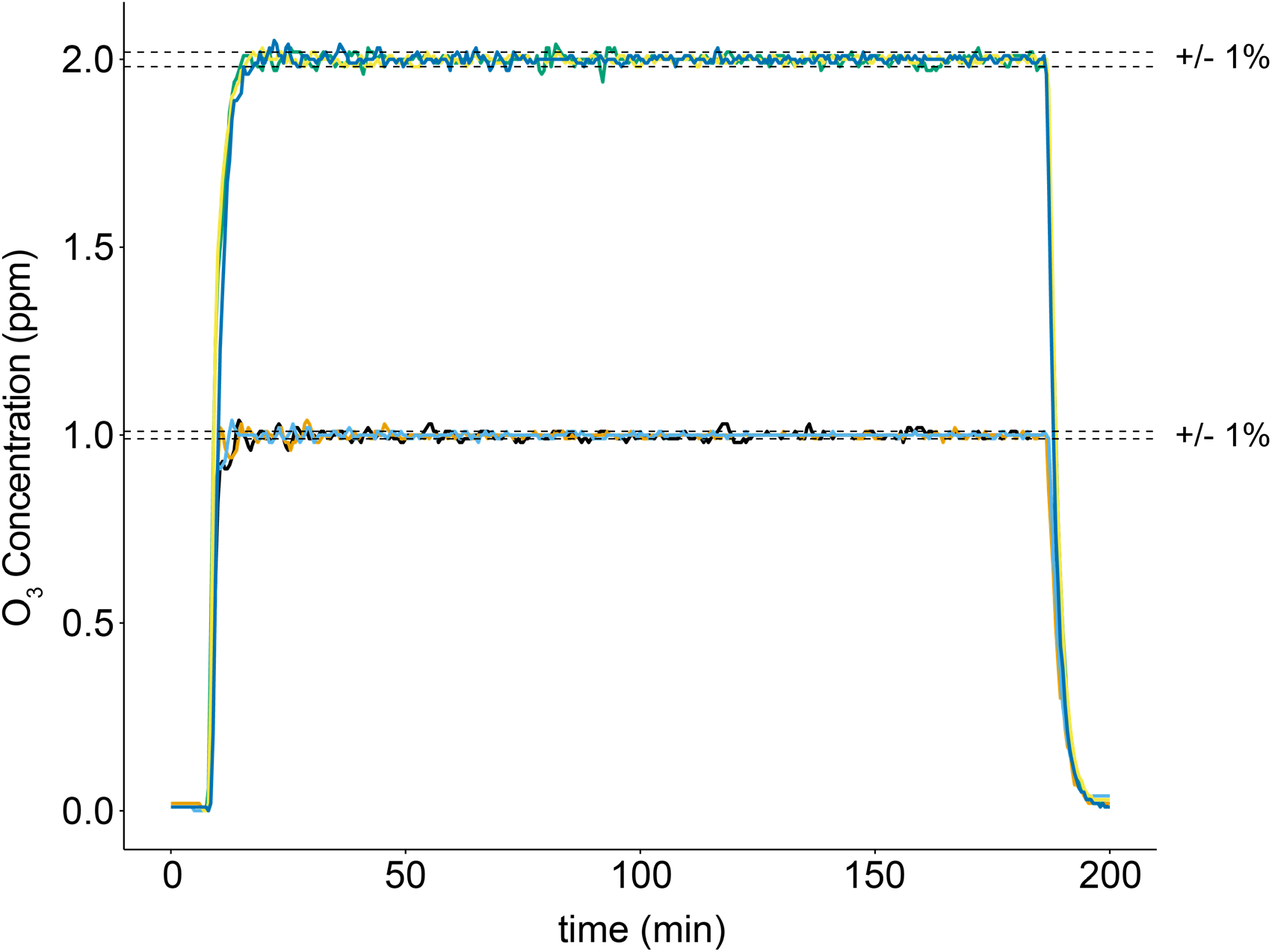
Concentration profiles for several exposures with mice present in the chamber. In exposure experiments conducted at both 1 and 2 ppm, the system maintained the O_3_ exposure concentration to within 1% of the setpoint for 3h. The number of mice for the exposure experiments shown range from 6 to 38. Plots represent concentration-time profiles (30-second running average) of three exposures at each concentration.

#### Concentration Uniformity

One of the objectives of this project was to support future studies using large numbers of genetically diverse mice. These studies require several exposure batches and variable spatial placement of mice of different strains within the chambers. Although the chamber design, ozone inlet, and airflow rate all contribute to mixing and spatial distribution of ozone in the chamber, perfect mixing is unlikely to be achieved. As such, we conducted an evaluation of ozone concentration across several chamber locations to obtain reference data for potential covariate adjustment or serve as the impetus to take measures to improve mixing. The concentration uniformity assay (as determined by the oxidation of BKI solution, see methods) revealed a mean absorbance of 1.149 across all indicator dishes for the ozone chamber and a CV of 14.6% (Table 2). The top rack had the highest mean absorbance value of 1.266 and largest CV of 16.8%. The middle and bottom racks had mean absorbance values of 1.143 and 1.038, and CVs of 7.6% and 7.2%, respectively.

**Table 2.** Concentration Uniformity Data.

| **Location** | **Mean Absorbance** | **SD** | **CV** |
| --- | --- | --- | --- |
| **Control (n=3)** | 0.034 | 0.004 | 13.098 |
| **Ozone** |  |  |  |
| Top Rack (n=5) | 1.266 | 0.212 | 16.768 |
| Middle Rack (n=5) | 1.143 | 0.087 | 7.639 |
| Bottom Rack (n=5) | 1.038 | 0.075 | 7.215 |
| **Ozone Overall (n=15)** | **1.149** | **0.168** | **14.594** |

*SD = Standard deviation.*
*CV = Coefficient of Variation (SD/Mean\*100)*

#### Animal Exposures

The biological consequences of acute ozone exposure include the induction of airway inflammation, which is in part reflected by the recruitment of neutrophils to the airways. To assess the reproducibility of this outcome in our exposure system, we measured the levels of airway neutrophilia across several experiments using the same strain (and sex) of mice. Data from four experiments in which female C57BL/6J mice were exposed to 2 ppm O_3_ or filtered air (Figure 5) demonstrate that exposures with our system produced biologically reproducible results.

**Figure 5.**
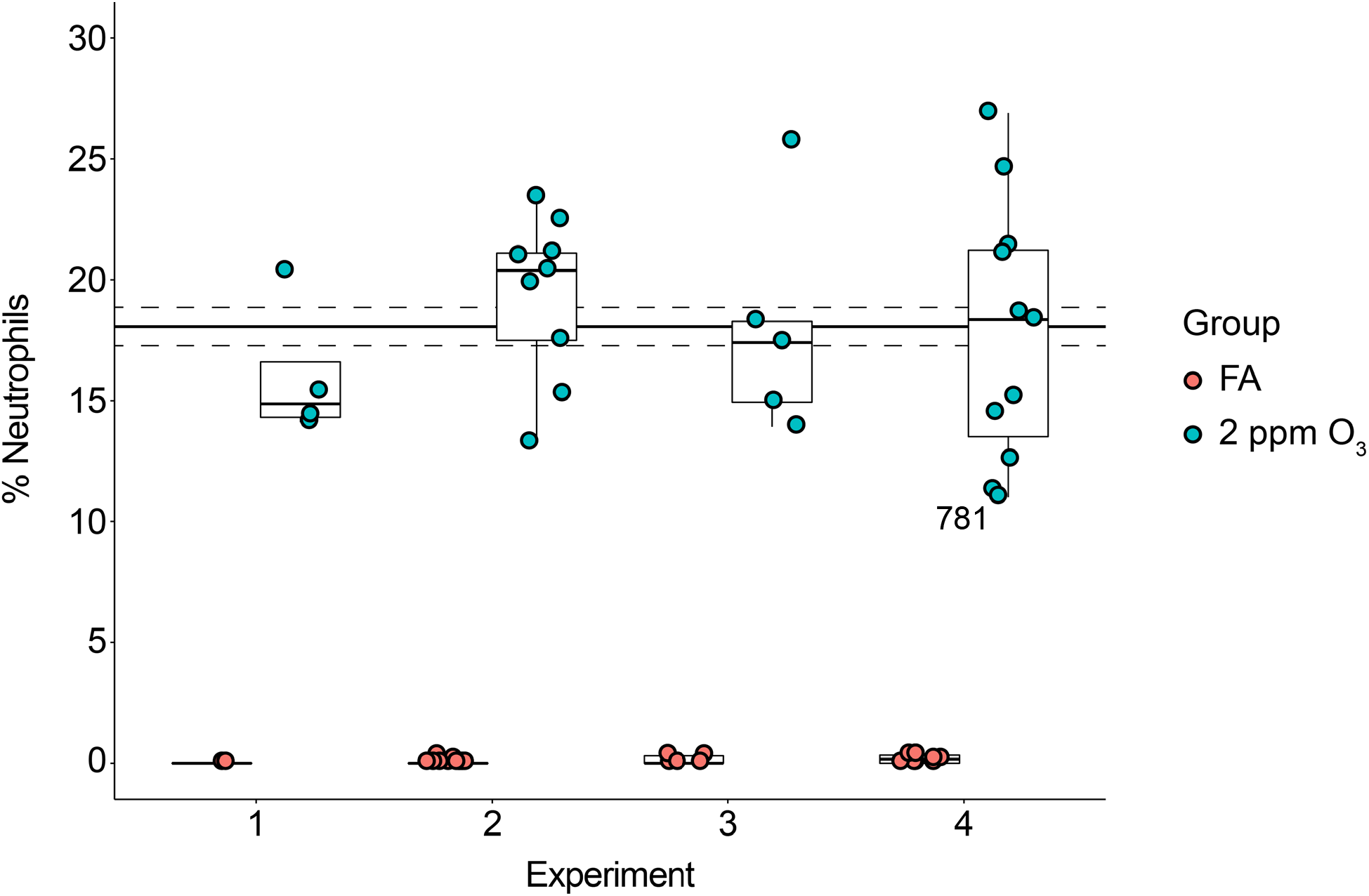
Elevated BAL fluid neutrophils in O_3_ exposed mice across four experiments. Experiments 1 through 4 were conducted over the span of several months to test the reproducibility of our exposure system by a biological measure. On average, the percent neutrophils was 18.1 ± 0.8% (solid and dashed lines, mean ± SEM), and the percent neutrophil value was not significantly different across experiments (p > 0.05, holm corrected pairwise t-test).

## Discussion

Our goal for this project was to develop a whole-body O_3_ inhalation exposure system, suitable for large numbers of mice. With the present design, we achieved a system that performed well and met our major requirements in terms of safety, capacity, reproducibility, automation, and data management. Our safety cutoff system operated correctly, turning off the flow of O_3_ upon the requisite increase in chamber static pressure. During experiments, the system was straightforward to setup, run, and monitor remotely. With the system under automated PID control, we were able to maintain consistent and repeatable exposure profiles with up to 38 mice present in the chambers. As expected, due to imperfect mixing we measured minor differences in the concentration of O_3_ (as reflected by the BKI assay) between fifteen chamber locations. The most variation was measured across the top rack of the chamber and the center dish of this rack had the highest absorbance overall. Although there was similar variation in the control absorbance values, it may be prudent to avoid this top-center region when placing mice in the chamber if possible. Nonetheless, in each experiment we conducted, ozone concentrations were maintained to within 1% of the desired concentration. Additionally, we measured consistent toxicologic responses to 2 ppm ozone over several exposures, confirming the system’s overall performance and repeatability.

We based the design of our system primarily on a much larger facility at the U.S. EPA’s Inhalation Toxicology Facilities Branch in RTP, NC. This facility houses a number of laboratory spaces with various styles of chambers and capabilities for a variety of air pollutant exposures. Several of the components of our system are the same as the EPA ozone exposure system, namely the H-1000 chambers, arrangement of the clean air supply and exhaust, DASYLab^®^ control software, and Thermo 49i O_3_ analyzer. Because of the relatively smaller size of our system (2 vs 4 chambers at the EPA) and laboratory, we made some decisions to use or construct different components. For example, unique and custom-built components of our system include the stainless steel supply ductwork, CPVC exhaust pipes, the arrangement and organization of manifolds and control valves, remote tubing to monitor pressures, and the power supply and analog signal cable wiring. Different commercially acquired components include the supply and exhaust fans, filtration unit, ozone and oxygen generators, USB DAQ, and mass flow controllers. Despite these differences, we were able to achieve a similar level of performance as the EPA system in terms of automated and reproducible exposures (Higuchi and Walsh, personal communications).

In comparison to published ozone exposure systems, ours is most similar to one developed by investigators McKinney and Frazer at NIOSH (McKinney and Frazer 2008). We employed a similar feedback loop for automated control of O_3_ as used by McKinney and Frazer. However, our chambers are substantially larger, and thus required different methods for generating both diluent air and ozone. Our diluent air is supplied by an electric pressure blower that pushes laboratory air through a series of filter panels as opposed to dry compressed air. We chose to use a corona arc discharge ozone generator because of the greater output capacity than a mercury lamp-style ozone generator. Additionally, compared to the NIOSH system, temperature and humidity in our chambers are controlled by the building HVAC system and we integrate environmental data recording (pressure, temperature, and relative humidity) alongside the feedback control of ozone exposure using DASYLab^®^. Using this arrangement, we were able to maintain chamber temperature and humidity during and across exposure experiments within the Office of Economic Cooperation and Development (OECD) recommended ranges of 22 ± 3°C and 30 to 70% relative humidity (OECD 2018).

While using the system over time, we identified a few areas for potential improvement or modifications. There are reports that changes in humidity can alter or interfere with the readings of UV absorbance ozone analyzers (Wilson and Birks 2006). Because the humidity did not change dramatically during or between experiments, we did not investigate whether water vapor interference was an issue for our system. In either case, installing lengths of Nafion tubing on both the sample and reference lines just prior to the photometer has been shown to eliminate any potential water vapor interference and thus may be a good practice. (Note: care should be taken to make sure calibrations are performed under the same conditions as typical sampling (Wilson and Birks 2006).) More precise and direct/local control of chamber humidity and temperature using an additional feedback control system may also help to address inter-experimental variability and water vapor interference caused by changes in humidity. While there is no standard for acceptable concentration uniformity and our data on chamber concentration uniformity from the BKI assay are within the range seen by other groups generating air pollutant exposures in H-1000 chambers (Marra and Rombout 1990), we may improve ozone concentration uniformity by increasing chamber airflow or installing a recirculation system. Finally, we observed that the temperature increases in the safety cabinet because of heat produced by the equipment therein. Excessive heat may decrease the output of the ozone generator or decrease the life of the equipment housed in the cabinet, therefore it may be desirable to examine this issue in detail in the future.

In conclusion, we have developed a robust, computer-controlled ozone exposure system that can be used to expose up to 180 mice per experiment. Our system provides precise and reproducible ozone exposures, maintaining the desired ozone concentration regardless of the number of subjects. The system’s data acquisition and control equipment affords minimal operator involvement during exposures and automatic recording of data on environmental conditions. This new system will be invaluable for large-scale studies in mice and facilitate the identification of mechanisms by which ozone causes adverse respiratory and systemic health effects.

## Supporting information

## Acknowledgements

We thank the following individuals who were instrumental to the success of this project: Ian Gilmour at The United States Environmental Protection Agency’s National Health and Environmental Effects Research Laboratory (NHEERL), Chris Gregory from the Department of Genetics at The University of North Carolina Chapel Hill, and William Robertson and Artie Neese of Facilities Services at The University of North Carolina Chapel Hill. This research was supported by the National Institute of Environmental Health Sciences under award numbers R01 ES024965, T32 ES007126-35, and P30-ES-010126.

## References

Arab K, Mp A. 2012. PID Control Theory. In: Introduction to PID Controllers - Theory, Tuning and Application to Frontier Areas.

Aris RM, Christian D, Hearne PQ, Kerr K, Finkbeiner WE, Balmes JR. 1993. Ozone-induced airway inflammation in human subjects as determined by airway lavage and biopsy. Am Rev Respir Dis. 148(5):1363–72. doi:10.1164/ajrccm/148.5.1363.

Astrom K, Hägglund T. 1995. PID controllers: theory, design and tuning. Instrum Soc Am. doi:1556175167.

Bauer AK, Kleeberger SR. 2010. Genetic mechanisms of susceptibility to ozone-induced lung disease. Ann N Y Acad Sci. 1203(1):113–119. doi:10.1111/j.1749-6632.2010.05606.x. [accessed 2018 Oct 23]. http://www.ncbi.nlm.nih.gov/pubmed/20716292.

Brown MG, Moss OR. 1981. An inhalation exposure chamber designed for animal handling. Lab Anim Sci. 31(6):717–20. [accessed 2018 Oct 18]. http://www.ncbi.nlm.nih.gov/pubmed/7343770.

Cabello N, Mishra V, Sinha U, DiAngelo SL, Chroneos ZC, Ekpa NA, Cooper TK, Caruso CR, Silveyra P. 2015. Sex differences in the expression of lung inflammatory mediators in response to ozone. Am J Physiol Lung Cell Mol Physiol. 309(10):L1150–63. doi:10.1152/ajplung.00018.2015.

Chen L-C, Lippmann M. 2015. Inhalation toxicology methods: the generation and characterization of exposure atmospheres and inhalational exposures. Curr Protoc Toxicol. 63:24.4.1–23. doi:10.1002/0471140856.tx2404s63. [accessed 2018 Sep 25]. http://www.ncbi.nlm.nih.gov/pubmed/25645246.

Cho Y, Abu-Ali G, Tashiro H, Brown TA, Osgood R, Kasahara DI, Huttenhower C, Shore SA. 2018 Sep 21. Sex Differences in Pulmonary Responses to Ozone in Mice: Role of the Microbiome. Am J Respir Cell Mol Biol.:rcmb.2018-0099OC. doi:10.1165/rcmb.2018-0099OC. [accessed 2018 Sep 25]. https://www.atsjournals.org/doi/10.1165/rcmb.2018-0099OC.

Cleary EG, Cifuentes M, Grinstein G, Brugge D, Shea TB. 2018. Association of Low-Level Ozone with Cognitive Decline in Older Adults. J Alzheimers Dis. 61(1):67–78. doi:10.3233/JAD-170658. [accessed 2018 Oct 23]. http://www.ncbi.nlm.nih.gov/pubmed/29103040.

Day DB, Xiang J, Mo J, Li F, Chung M, Gong J, Weschler CJ, Ohman-Strickland PA, Sundell J, Weng W, et al. 2017. Association of Ozone Exposure With Cardiorespiratory Pathophysiologic Mechanisms in Healthy Adults. JAMA Intern Med. 177(9):1344. doi:10.1001/jamainternmed.2017.2842. [accessed 2018 Sep 25]. http://www.ncbi.nlm.nih.gov/pubmed/28715576.

Devlin RB, McDonnell WF, Becker S, Madden MC, McGee MP, Perez R, Hatch G, House DE, Koren HS. 1996. Time-dependent changes of inflammatory mediators in the lungs of humans exposed to 0.4ppm ozone for 2hr: A comparison of mediators found in broncoalveolar lavage fluid 1 and 18 hr after exposure. Toxicol Appl Pharmacol. 138:176–85.

Dorato MA. 1990. Overview of inhalation toxicology. Environ Health Perspect. 85:163–170. doi:10.2307/3430680.

Goldsmith WT, McKinney W, Jackson M, Law B, Bledsoe T, Siegel P, Cumpston J, Frazer D. 2011. A computer-controlled whole-body inhalation exposure system for the oil dispersant COREXIT EC9500A. J Toxicol Environ Health A. 74(21):1368–80. doi:10.1080/15287394.2011.606793. [accessed 2018 Oct 17]. http://www.ncbi.nlm.nih.gov/pubmed/21916743.

Herring MJ, Putney LF, St George JA, Avdalovic M V, Schelegle ES, Miller LA, Hyde DM. 2015. Early life exposure to allergen and ozone results in altered development in adolescent rhesus macaque lungs. Toxicol Appl Pharmacol. 283(1):35–41. doi:10.1016/j.taap.2014.12.006. [accessed 2018 Oct 18]. https://linkinghub.elsevier.com/retrieve/pii/S0041008X1400444X.

Ito K, De Leon SF, Lippmann M. 2005. Associations between ozone and daily mortality: Analysis and meta-analysis. Epidemiology. 16(4):446–457. doi:10.1097/01.ede.0000165821.90114.7f.

Jerrett M, Brook R, White LF, Burnett RT, Yu J, Su J, Seto E, Marshall J, Palmer JR, Rosenberg L, et al. 2017. Ambient ozone and incident diabetes: A prospective analysis in a large cohort of African American women. Environ Int. 102:42–47. doi:10.1016/j.envint.2016.12.011. [accessed 2018 Sep 25]. https://linkinghub.elsevier.com/retrieve/pii/S0160412016310078.

Jerrett M, Burnett RT, Pope CA, Ito K, Thurston G, Krewski D, Shi Y, Calle E, Thun M. 2009. Long-Term Ozone Exposure and Mortality. N Engl J Med. 360(11):1085–1095. doi:10.1056/NEJMoa0803894.

Ji M, Cohan DS, Bell ML. 2011. Meta-analysis of the Association between Short-Term Exposure to Ambient Ozone and Respiratory Hospital Admissions. Environ Res Lett. 6(2). doi:10.1088/1748-9326/6/2/024006. [accessed 2018 Sep 25]. http://www.ncbi.nlm.nih.gov/pubmed/21779304.

Kim B-J, Kwon J-W, Seo J-H, Kim H-B, Lee S-Y, Park K-S, Yu J, Kim H-C, Leem J-H, Sakong J, et al. 2011. Association of ozone exposure with asthma, allergic rhinitis, and allergic sensitization. Ann Allergy, Asthma Immunol. 107(3):214–219.e1. doi:10.1016/j.anai.2011.05.025. [accessed 2017 Mar 1]. http://linkinghub.elsevier.com/retrieve/pii/S1081120611003929.

Ko FWS, Tam W, Wong TW, Chan DPS, Tung AH, Lai CKW, Hui DSC. 2007. Temporal relationship between air pollutants and hospital admissions for chronic obstructive pulmonary disease in Hong Kong. Thorax. 62(9):780–5. doi:10.1136/thx.2006.076166. [accessed 2018 Sep 25]. http://www.ncbi.nlm.nih.gov/pubmed/17311838.

Levy JI, Carrothers TJ, Tuomisto JT, Hammitt JK, Evans JS. 2001. Assessing the public health benefits of reduced ozone concentrations. Environ Health Perspect. 109(12):1215–1226. doi:10.1289/ehp.011091215.

Levy JI, Chemerynski SM, Sarnat JA. 2005. Ozone exposure and mortality: An empiric bayes metaregression analysis. Epidemiology. 16(4):458–468. doi:10.1097/01.ede.0000165820.08301.b3.

Marra M, Rombout PJA. 1990. Design and Performance of an Inhalation Chamber for Exposing Laboratory Animals to Oxidant Air Pollutants. Inhal Toxicol. 2(3):187–204. doi:10.3109/08958379009145254.

McKinney W, Frazer D. 2008. Computer-controlled ozone inhalation exposure system. Inhal Toxicol. 20(1):43–8. doi:10.1080/08958370701758544.

Michaudel C, Fauconnier L, Julé Y, Ryffel B. 2018. Functional and morphological differences of the lung upon acute and chronic ozone exposure in mice. Sci Rep. 8(1):10611. doi:10.1038/s41598-018-28261-9. [accessed 2018 Oct 18]. http://www.ncbi.nlm.nih.gov/pubmed/30006538.

Miller DB, Ghio AJ, Karoly ED, Bell LN, Snow SJ, Madden MC, Soukup J, Cascio WE, Ian Gilmour M, Kodavanti UP. 2016. Ozone exposure increases circulating stress hormones and lipid metabolites in humans. Am J Respir Crit Care Med. 193(12):1382–1391. doi:10.1164/rccm.201508-1599OC.

Miller DB, Snow SJ, Schladweiler MC, Richards JE, Ghio AJ, Ledbetter AD, Kodavanti UP. 2016. Acute Ozone-Induced Pulmonary and Systemic Metabolic Effects Are Diminished in Adrenalectomized Rats. Toxicol Sci. 150(2):312–322. doi:10.1093/toxsci/kfv331.

Mudway IS, Kelly FJ. 2000. Ozone and the lung: A sensitive issue. Mol Aspects Med. 21(1–2):1–48. doi:10.1016/S0098-2997(00)00003-0.

Nielsen GD, Hougaard KS, Larsen ST, Hammer M, Wolkoff P, Clausen PA, Wilkins CK, Alarie Y. 1999. Acute airway effects of formaldehyde and ozone in BALB/c mice. Hum Exp Toxicol. 18(6):400–409. doi:10.1191/096032799678840246. [accessed 2017 Jan 29]. http://www.ncbi.nlm.nih.gov/pubmed/10413245.

O’Shaughnessy PT, Achutan C, O’Neill ME, Thorne PS. 2003. A small whole-body exposure chamber for laboratory use. Inhal Toxicol. 15(3):251–263. doi:10.1080/08958370304504\rU1HB4TQL7UH2FQWW [pii].

OECD. 2018. Acute Inhalation Toxicity: Fixed Concentration Procedure. (June). doi:10.1787/9789264284166-en. [accessed 2018 Oct 24]. https://www.oecd-ilibrary.org/environment/test-no-433-acute-inhalation-toxicity-fixed-concentration-procedure_9789264284166-en.

Paffett ML, Zychowski KE, Sheppard L, Robertson S, Weaver JM, Lucas SN, Campen MJ. 2015. Ozone Inhalation Impairs Coronary Artery Dilation via Intracellular Oxidative Stress: Evidence for Serum-Borne Factors as Drivers of Systemic Toxicity. Toxicol Sci. 146(2):244–53. doi:10.1093/toxsci/kfv093. [accessed 2018 Sep 25]. http://www.ncbi.nlm.nih.gov/pubmed/25962394.

Pauluhn J. 2003. Overview of testing methods used in inhalation toxicity: From facts to artifacts. Toxicol Lett. 140–141:183–193. doi:10.1016/S0378-4274(02)00509-X.

Phalen RF. 1976. Inhalation exposure of animals. Environ Health Perspect. Vol.16(August):17–24. doi:10.1289/ehp.761617.

Pryor WA, Squadrito GL, Friedman M. 1995. The cascade mechanism to explain ozone toxicity: The role of lipid ozonation products. Free Radic Biol Med. 19(6):935–941. doi:10.1016/0891-5849(95)02033-7.

Speen AM, Kim HH, Bauer RN, Meyer M, Gowdy KM, Fessler MB, Duncan KE, Liu W, Porter NA, Jaspers I. 2016. Ozone-derived Oxysterols Affect Liver X Receptor (LXR) Signaling: A Potential Role for Lipid-Protein Adducts. (2). doi:10.1074/jbc.M116.732362.

Turner MC, Jerrett M, Pope CA, Krewski D, Gapstur SM, Diver WR, Beckerman BS, Marshall JD, Su J, Crouse DL, et al. 2016. Long-Term Ozone Exposure and Mortality in a Large Prospective Study. Am J Respir Crit Care Med. 193(10):1134–1142. doi:10.1164/rccm.201508-1633OC. [accessed 2018 Sep 25]. http://www.atsjournals.org/doi/10.1164/rccm.201508-1633OC.

Tyler CR, Noor S, Young T, Rivero V, Sanchez B, Lucas S, Caldwell KK, Milligan ED, Campen MJ. 2018. Aging Exacerbates Neuroinflammatory Outcomes Induced by Acute Ozone Exposure. Toxicol Sci. doi:10.1093/toxsci/kfy014.

Wilson KL, Birks JW. 2006. Mechanism and elimination of a water vapor interference in the measurement of ozone by UV absorbance. Environ Sci Technol. 40(20):6361–6367. doi:10.1021/es052590c.

Wong BA. 2007. Inhalation Exposure Systems: Design, Methods and Operation. Toxicol Pathol. 35(1):3–14. doi:10.1080/01926230601060017. [accessed 2017 Mar 4]. http://www.ncbi.nlm.nih.gov/pubmed/17325967.

Zu K, Shi L, Prueitt RL, Liu X, Goodman JE. 2018. Critical review of long-term ozone exposure and asthma development. Inhal Toxicol. 30(3):99–113. doi:10.1080/08958378.2018.1455772. [accessed 2018 Oct 18]. https://www.tandfonline.com/doi/full/10.1080/08958378.2018.1455772.

