## Supplementary material for "Development of a large-scale computer-controlled ozone inhalation exposure system for rodents"

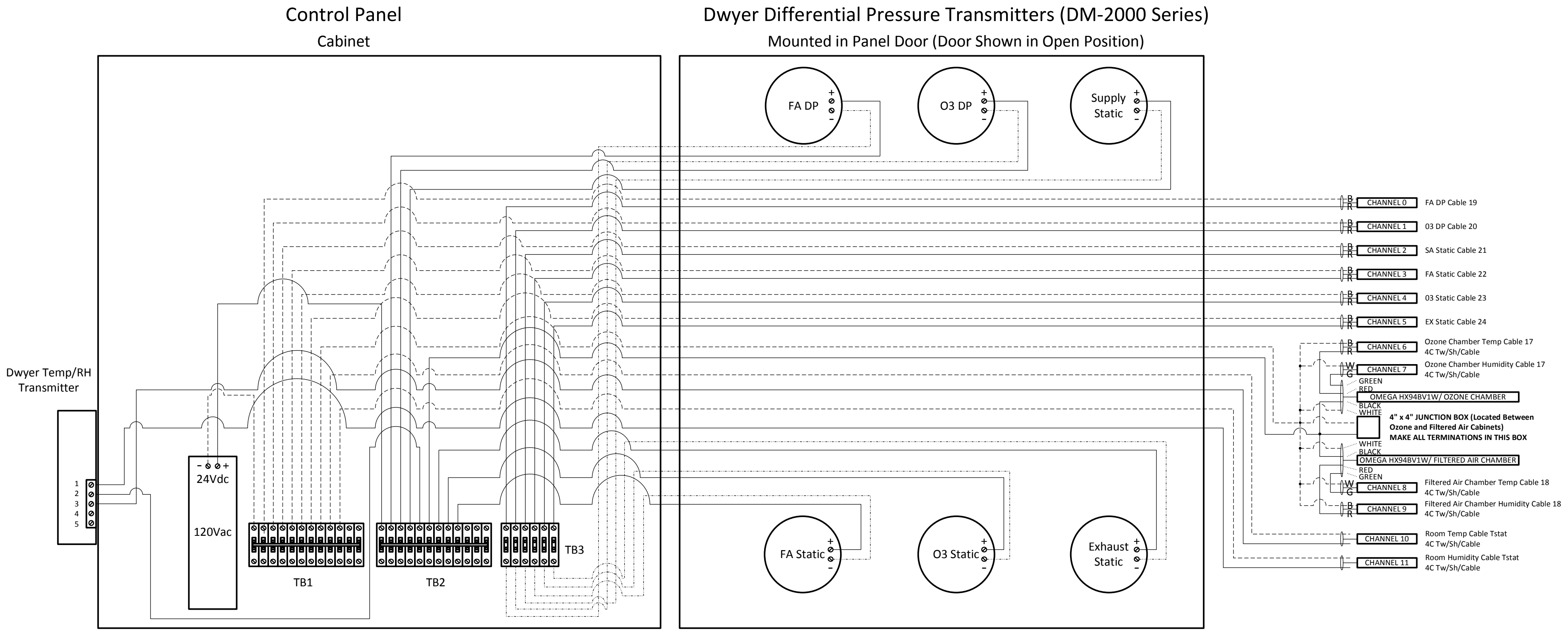


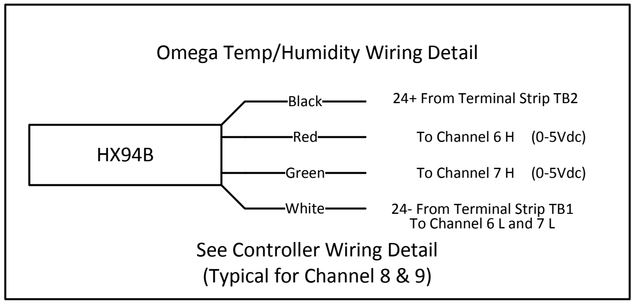


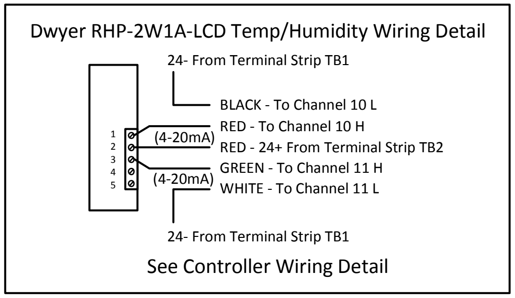


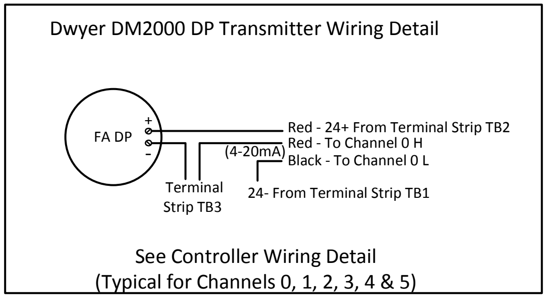


### Wall Cabinet Wiring

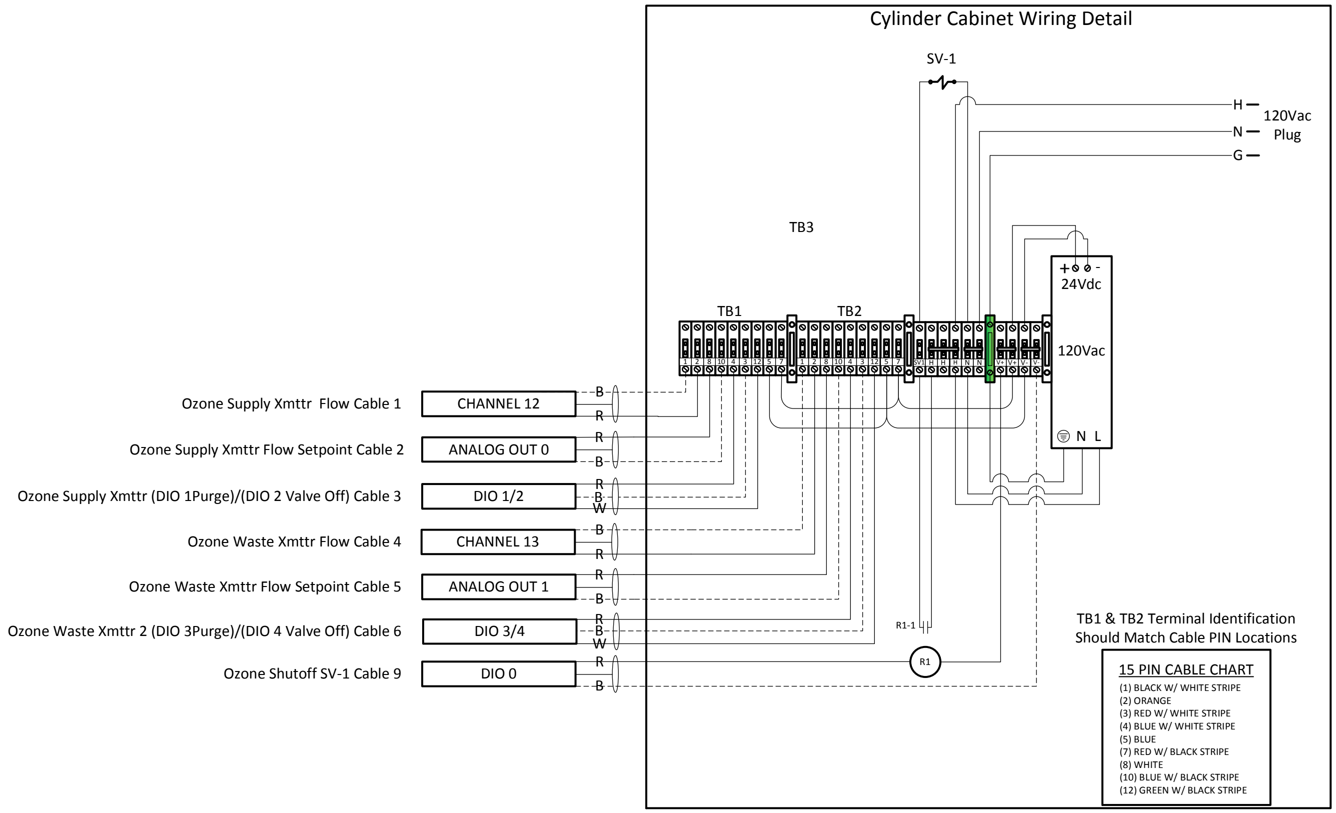


| PIN # | Function |
|---|---|
| 1 | 0-5Vdc Flow Signal Common |
| 2 | 0-5Vdc Flow Signal Output |
| 3 | Common |
| 4 | Purge |
| 5 | Power Supply Common |
| 7 | Power Supply +24Vdc |
| 8 | Remote Setpoint Input |
| 10 | Remote Setpoint Common |
| 12 | Valve Off Control |
| 1 & 2 | 0-5Vdc Flow Signal Output |
| 3 & 4 | Purge |
| 3 & 12 | Valve Off Control |
| 5 & 7 | 24 Vdc Power Supply to Mass Flow Controller |
| 8 & 10 | 0-5Vdc Remote Setpoint |

### Gas Cylinder Cabinet Wiring

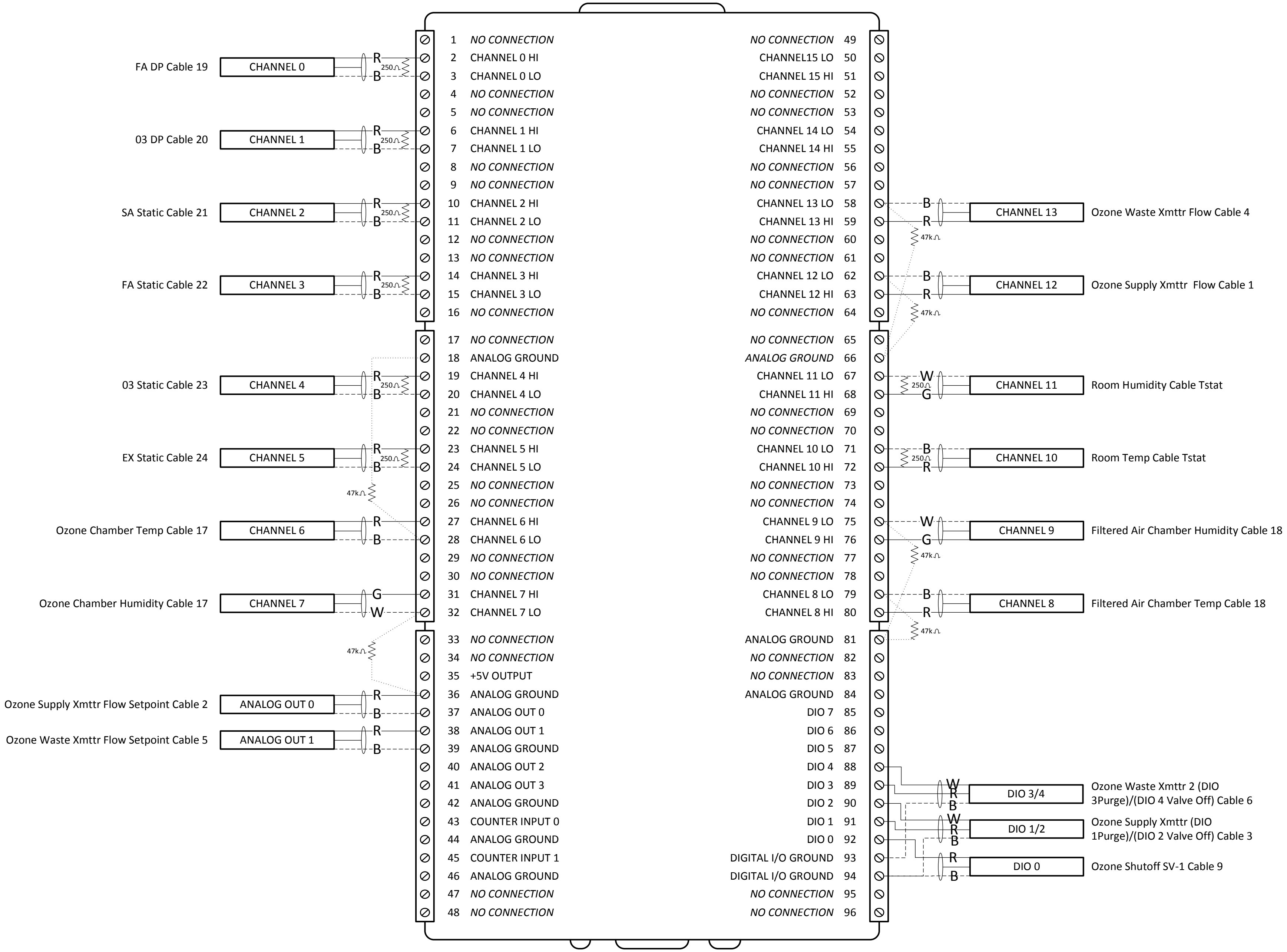


### USB-2416-4AO Multi Functional DAQ Device Wiring

| Point Address | Point Type | Point Description | Cable # | Wire Color | Further Description |
|---|---|---|---|---|---|
| DO0 | Digital Output | Solenoid Valve SV-1 | 9 | Red | Relay 5 in Cylinder Cabinet/120V Omega Solenoid |
| Ground | | | | Black | Connected to Relay 5 Common |
| DO1 | Digital Output | Mass Flow Xmttr 1 Purge | 3 | Red | Relay 1 in Cylinder Cabinet/Connected to PIN 4 AALBORG 15 PIN Cable |
| Ground | | | | Black | Relay 1 & 2 in Cylinder Cabinet |
| DO2 | Digital Output | Mass Flow Xmttr 1 Valve Off | | White | Relay 2 in Cylinder Cabinet/Connected to PIN 12 AALBORG 15 PIN Cable |
| DO3 | Digital Output | Mass Flow Xmttr 2 Purge | 6 | Red | Relay 3 in Cylinder Cabinet/Connected to PIN 4 AALBORG 15 PIN Cable |
| Ground | | | | Black | Relay 3 & 4 in Cylinder Cabinet |
| DO4 | Digital Output | Mass Flow Xmttr 2 Valve Off | | White | Relay 4 in Cylinder Cabinet/Connected to PIN 12 AALBORG 15 PIN Cable |
| AO0 | 0-5Vdc | Mass Flow Xmttr 1 Flow Setpoint | 2 | Red | Connected to PIN 8 AALBORG 15 PIN Cable |
| Ground | | | | Black | Connected to PIN 10 AALBORG 15 PIN Cable |
| AO1 | 0-5Vdc | Mass Flow Xmttr 2 Flow Setpoint | 5 | Red | Connected to PIN 8 AALBORG 15 PIN Cable |
| Ground | | | | Black | Connected to PIN 10 AALBORG 15 PIN Cable |
| Channel 0 Hi | 4-20mA* | Filtered Air Chamber Differential | 19 | Red | COM on Dwyer DP Xmttr |
| Channel 0 Lo | | | | Black | Power Supply Ground |
| Channel 1 Hi | 4-20mA* | Ozone Chamber Differential | 20 | Red | COM on Dwyer DP Xmttr |
| Channel 1 Lo | | | | Black | Power Supply Ground |
| Channel 2 Hi | 4-20mA* | Supply Air Static | 21 | Red | COM on Dwyer DP Xmttr |
| Channel 2 Lo | | | | Black | Power Supply Ground |
| Channel 3 Hi | 4-20mA* | Filtered Air Chamber Static | 22 | Red | COM on Dwyer DP Xmttr |
| Channel3 Lo | | | | Black | Power Supply Ground |
| Channel 4 Hi | 4-20mA* | Ozone Cabinet Static | 23 | Red | COM on Dwyer DP Xmttr |
| Channel 4 Lo | | | | Black | Power Supply Ground |
| Channel5 Hi | 4-20mA* | Exhaust Static | 24 | Red | COM on Dwyer DP Xmttr |
| Channel 5 Lo | | | | Black | Power Supply Ground |
| Channel 6 Hi | 0-5Vdc** | Ozone Chamber Temperature | 17 | Red | Connected to Red Lead Wire on OMEGA HX94B |
| Channel 6 Lo | | Common | | Black | Connected to White Lead Wire on OMEGA HX94B |
| Channel7 Hi | 0-5Vdc** | Ozone Chamber Humidity | | Green | Connected to Green Lead Wire on OMEGA HX94B |
| Channel7 Lo | | Common | | White | Connected to White Lead Wire on OMEGA HX94B |
| Channel 8 Hi | 0-5Vdc** | Filtered Air Chamber Temperature | 18 | Red | Connected to Red Lead Wire on OMEGA HX94B |
| Channel 8 Lo | | Common | | Black | Connected to White Lead Wire on OMEGA HX94B |
| Channel 9 Hi | 0-5Vdc** | Filtered Air Chamber Humidity | | Green | Connected to Green Lead Wire on OMEGA HX94B |
| Channel 9 Lo | | Common | | White | Connected to White Lead Wire on OMEGA HX94B |
| Channel 10 Hi | 4-20mA* | Room Temperature | T'Stat | Red | Connected to Terminal 1 on Dwyer RHP-2W1X-LCD Xmttr |
| Channel 10 Lo | | Common | | Black | Connected to TB1 in Wall Cabinet |
| Channel 11 Hi | 4-20mA* | Room Humidity | | Green | Connected to Terminal 3 on Dwyer RHP-2W1X-LCD Xmttr |
| Channel 11 Lo | | Common | | White | Connected to TB1 in Wall Cabinet |
| Channel 12 Hi | 0-5Vdc** | Mass Flow Xmttr 1 Flow | 1 | Red | Connected to PIN 2 AALBORG 15 PIN Cable |
| Channel 12 Lo | | Common | | Black | Connected to PIN 1 AALBORG 15 PIN Cable |
| Channel 13 Hi | 0-5Vdc** | Mass Flow Xmttr 2 Flow | 4 | Red | Connected to PIN 2 AALBORG 15 PIN Cable |
| Channel 13 Lo | | Common | | Black | Connected to PIN 1 AALBORG 15 PIN Cable |
| Channel 14 Hi | | Not Used | | | |
| Channel 14 Lo | | Not Used | | | |
| Channel 15 Hi | | Not Used | | | |
| Channel 15 Lo | | Not Used | | | |

* 4-20mA Inputs require a 250 Ohm resistor to be wired from Channelx Hi and Lo terminals of the USB-2416-4AO DAQ Device.

** 0-5Vdc Inputs require a 47k Ohm resistor to be wired from Channelx Lo to Analog Ground on the USB-2416-4AO DAQ Device.

### USB-2416-4AO Multi Functional DAQ Device Wiring Detail
